# Efficient estimation of evolutionary rates by covariance aware regression

**DOI:** 10.1101/408005

**Authors:** Richard A. Neher

## Abstract

Shared ancestry among individuals results in correlated traits and these dependencies need to be accounted for in probabilistic inference. In strictly asexual populations, the covariances have a particularly simple block-like structure imposed by the phylogenetic tree. Ho and Ane showed how this block-like structure can be exploited to efficiently invert covariance matrices and fit linear models on trees in linear time. In this short note, I use these methods to estimate evolutionary rates and to find the root of the tree that optimizes the time-divergence relationship. The algorithm is implemented in TreeTime and can be used to estimate evolutionary rates and their confidence intervals with computational cost scaling linearly in the number of tips.

---

Individuals related by a phylogenetic tree and share ancestral lineages to various degrees: All individuals share the history prior to the most recent common ancestor (MRCA) and each split in the tree partitions the populations into groups that subsequently evolve independently along different paths. Any heritable property that changes through time along branches of the tree is therefore correlated among individuals. Closely related individuals tend to have similar properties and biggest differences will be found among individuals whose MRCA lived far in the past. Such phylogenetic correlations are common confounders in inference from sequence data, for example in contact map prediction from covarying amino acids (Dunn *et al.*, 2008), genome wide association studies (Price *et al.*, 2010), or in the analysis of quantitative traits in comparative phylogenetics (Freckleton, 2012). Correction of population structure and phylogenetic correlations generically requires inversion of the covariance matrix of the trait among individuals in the population.

In general, matrix inversion is a computationally expensive operation, but (Ho *et al.*, 2014) have shown that tree-structured covariance matrices can be inverted recursively. Furthermore, linear models don’t require matrix inversion but can be fit in linear time by recursively calculating weighted moments that account for covariation. The purpose of this note is to give a simplified albeit less general derivation of this algorithm and use it to estimate evolutionary rates from serially sampled sequence data.

Divergence from the root of the tree, i.e. the number of mutations per genome length *L* that accumulated in the sampled individuals since the MRCA of the entire sample (aka root-to-tip distance, *d*), is a simple example of a property with strong phylogenetic correlation. According to the molecular clock hypothesis, one expects *d*_*i*_ to increase linearly in time as *d*_*i*_ *≈* β(*t*_*i*_ *-T*_*M RCA*_), where *β* is the evolutionary rate (Zuckerkandl and Pauling, 1965). The actual values *d*_*i*_ of a specific tip *i* will fluctuate around the expectation β(*t*_*i*_ *-T*_*M RCA*_) since mutations accumulate stochastically, e.g. according to a Poisson process. Divergences of closely related tips are strongly correlated because they share most of their history, while divergences of tips that only merge at the root are independent. The root-to-tip distances are sums of independent contributions of branches *p*_*i*_ in the tree connecting the tip to the root. The mean and variance of divergence *d*_*i*_ can be expressed as

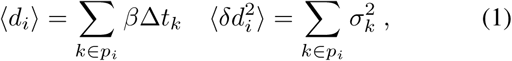

where Δ*t*_*k*_ is the length of the branch in calendar time and 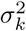 is the variance in divergence on a branch of length Δ*t*_*k*_ (in the simplest Poisson model 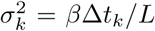). Divergences of different tips *i* of the tree, however, are clearly not independent observations and the covariance of *d*_*i*_ and *d*_*j*_ is given by

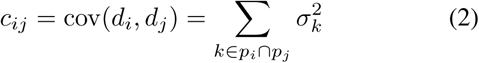

where the sum runs over all branches *k* in the intersection *p*_*i*_ ∩ *p*_*j*_ of the paths to the root of tips *i* and *j*. Fig. 1 shows this covariance matrix for a small example. The covariance matrix has a block like structure where each block corresponds to a branch in the tree that adds an identical contribution to all pairs of its descendants. This property naturally generates the three-point structure defined by Ho *et al.* (2014).

**FIG. 1.**
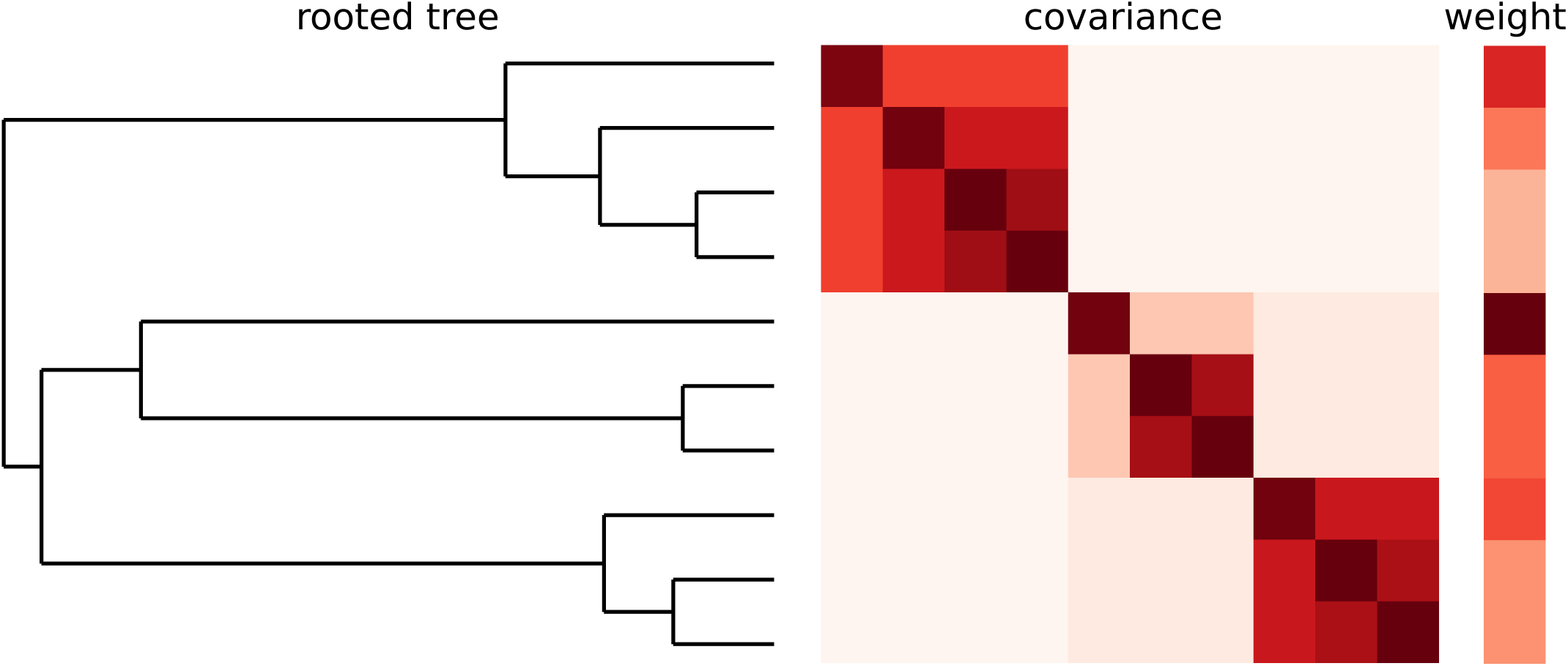
Covariance induced by a phylogenetic tree. The closer two tips are in the tree (left), the stronger are correlations among quantities that evolve along the tree (middle, darker colors). In an inference problem, tips need to be down-weighted if they are strongly correlated with other tips. The weight of a tip *i* is denoted by *r*_*i*_ in the text and is shown in the column on the right. The tip with the longest terminal branch has the largest weight *r*_*i*_.

This covariation complicates inference from tree-structured data. A common objective is to estimate the evolutionary rate *β* and *T*_*M RCA*_ of measurably evolving populations such as rapidly evolving RNA viruses (Drummond *et al.*, 2003b). The simplest approach would be to regress the root-to-tip distances *d*_*i*_ against sampling data *t*_*i*_ of all available tips by minimizing the squared deviation

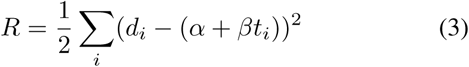

where *α* = *-β/T*_*M RCA*_ (Drummond *et al.*, 2003a; Paradis *et al.*, 2004; Rambaut *et al.*, 2016; Sagulenko *et al.*, 2018; To *et al.*, 2016; Volz and Frost, 2017). However, it is obvious that different data points (tips of the tree) are not independent observations and that simple least-squares regression will give noisy estimates of the evolutionary rate *β* or the *T*_*M RCA*_ with-out meaningful confidence intervals.

Instead, the *d*_*i*_ from all nodes are drawn approximately from a multivariate Gaussian distribution with covariance matrix **C** given by Eq. (2). An approximately most likely clock model should therefore minimize

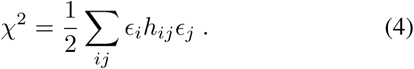

where *h*_*ij*_, **H** is the inverse of the covariance matrix **C** and ϵ_*i*_ = *d*_*i*_ *-α - t*_*i*_ *β* are the residuals. (Note that this ignores the prefactor involving determinant of **C**, but this prefactor is insignificant for sufficiently large dat sets.) Solving for *α,β* that minimize *χ*^2^ is straightforward but requires inversion of a potentially large covariance matrix which is a computationally expensive operation. Most implementations for matrix inversion scale as *𝒪*(*n*^3^) with the rank *n* of the matrix. Furthermore, the position of the root is typically unknown and the optimal model parameters have to be determined along with an optimization of the root. Naively, this would require repeating matrix inversion and model estimation for many choices of the root. The typical approach to circumvent this problem is to estimate to model and tree topology jointly by computationally expensive sampling of parameter and tree space (Drummond *et al.*, 2012).

As Ho *et al.* (2014) have shown, covariance matrices that have the tree-structure defined in Eq. (2) can be inverted recursively in *𝒪*(*n*^2^) operations. Furthermore, model parameters that minimize *χ*^2^ can be obtained in *𝒪*(*n*) operations without ever inverting the full matrix. And the recursive nature of the problem allows to evaluate the optimal model parameters for every possible choice of the root simultaneously without any additional computational burden. I will first rederive the recursive matrix inversion for a specific choice of root, then show how this calculation be efficiently done for every possible choice of root, and finally show how *χ*^2^ can be minimized without explicit calculation of the inverse covariance matrix for a linear model.

## Tree-covariance matrix inversion

Due to the structure of the covariance in Eq. (2) induced by the tree, **C** can be inverted recursively (Ho *et al.*, 2014). Consider first only the correlation **C**_*p*_ between leaves of node *p* induced by the child-subtrees of node *p*, see Fig. 2. Relative to node *p*, leaves that descend from the different child branches of *p* are uncorrelated and **C**_*p*_ has two (or more, one for each child) blocks on the diagonal. These blocks are sum of the analogous partial correlation matrices 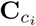 of the children *c*_*i*_ and the variation 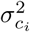 associated with the branch leading to child *i*:

**FIG. 2.**
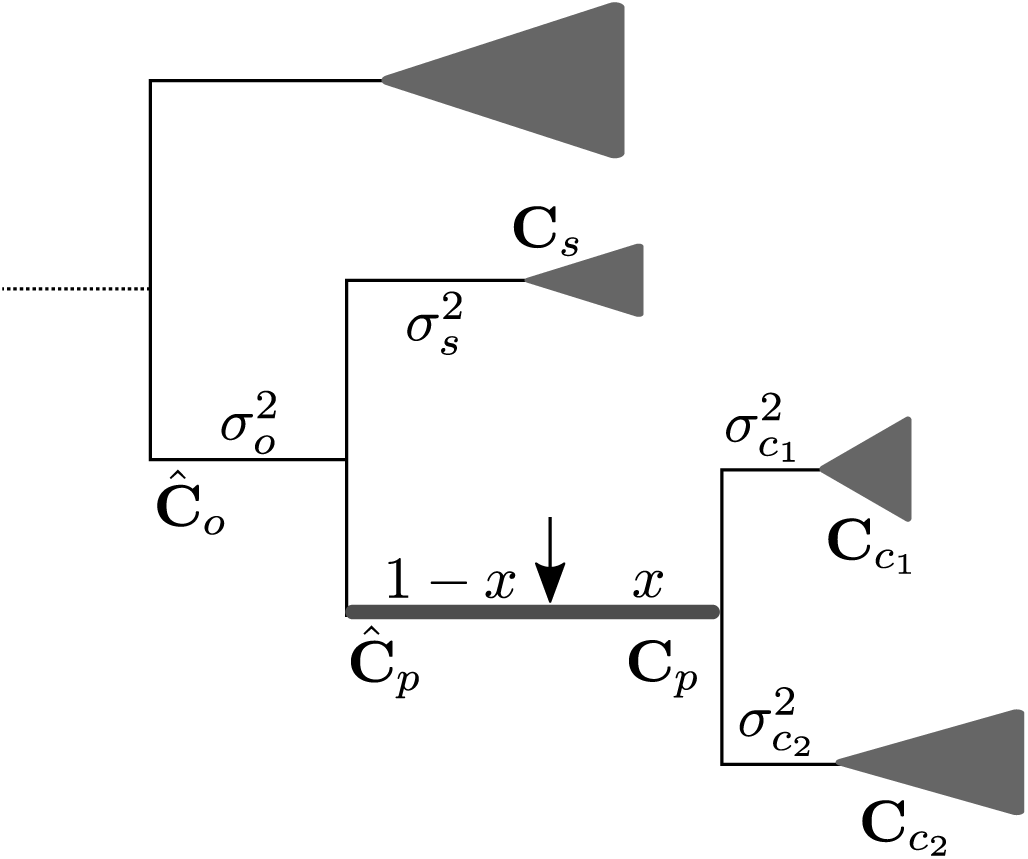
Illustration of the recursive inversion algorithm. Each internal node *p* is dressed with a covariance matrix **C**_*p*_ and its inverse **H**_*p*_ (not shown) of divergence of its leaves relative to *p*. These matrices can be calculated recursively from the analogous quantities of the children and the branch variances 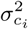, see Eq. (5) and Eq. (7). The covariance matrix of all leaves in the outgroup can be calculated in an analogous manner. Instead of the child nodes, this matrix includes one block for each sister clade *s* and one block for the outgroup of the parent node. To calculate the total leaf-covariance (or its inverse) for a choice of root – indicated by an arrow – at position *x* along the branch *p*, **C**_*p*_ and 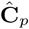 (or **H**_*p*_ and 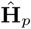) have to be combined in one matrix with additional variance 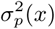 and 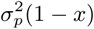.

**FIG. 3.**
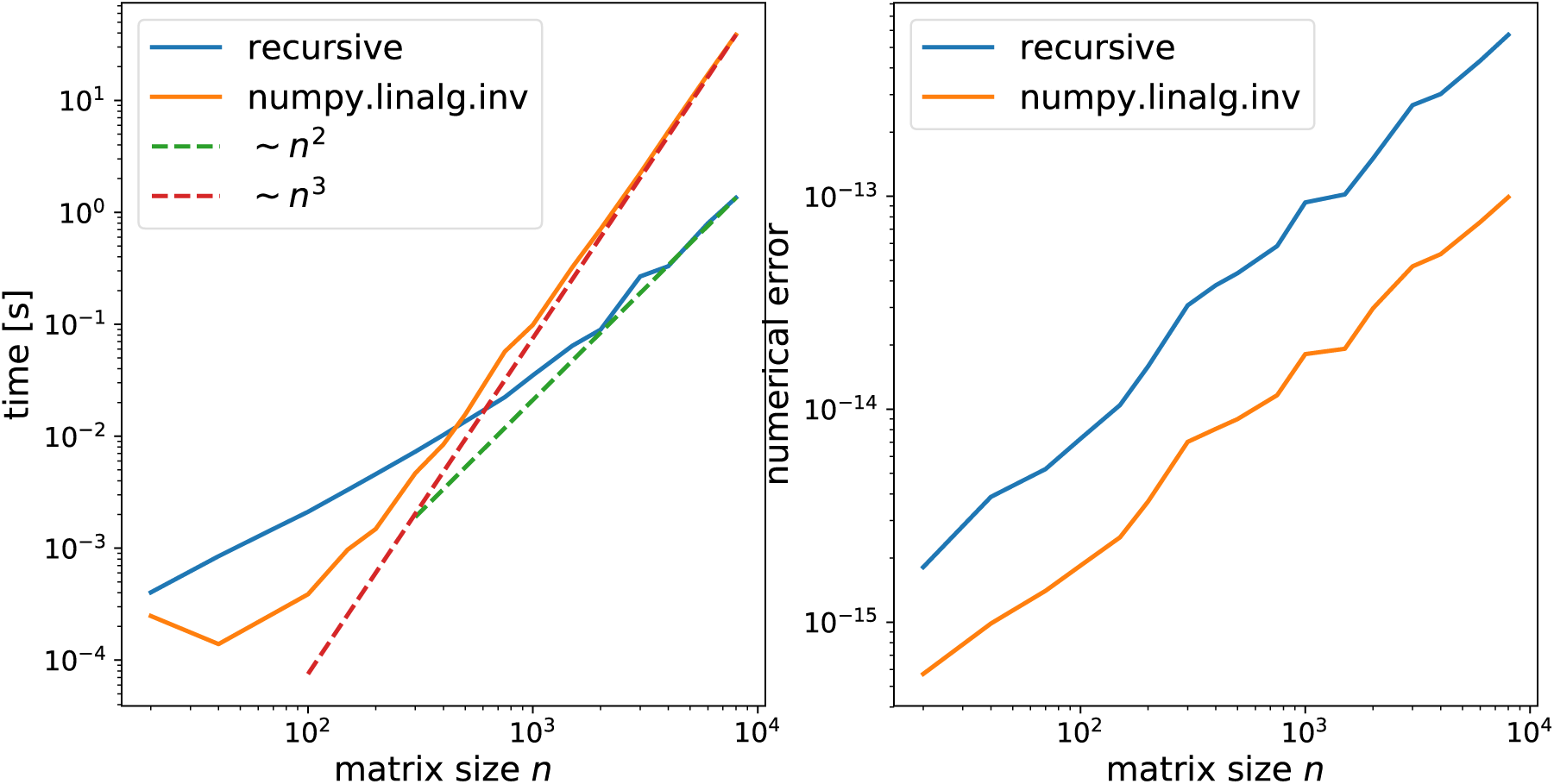
Speed and accuracy: While standard matrix inversion scales as *n*^3^ with rank of the matrix, the recursive algorithm scales much more favorably. In fact, up to n = 1000 the simply python/numpy implementation is dominated by tree-traversal and allocation and has not reached the asymptotic ∼ *n*^2^ scaling yet. The right panel shows the numerical accuracy of matrix inversion quantified as *n*^-2^ ∑_ij_ |δ_*ij*_ – ∑_*k*_ *c*_*ik*_*h*_*kj*_|. The recursive inverse is less accurate than `numpy.linalg.inv` but behaves similarly when the rank of the matrix changes.

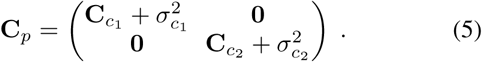

Inverting this matrix is cheap for two reasons: (i) the matrix is block diagonal, hence its inverse is the block diagonal of the inverse of its blocks. (ii) the individual blocks are a sum of a matrix with known inverse (calculated in the analogous step for child nodes) and a constant. The inverse of a the sum of a matrix with known inverse and constant can be calculated using the Sherman-Morrison-Woodbury Formula (Hager, 1989):

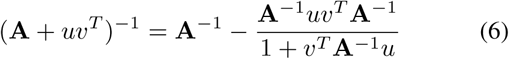

In our case, the outer product *uv*^*T*^ is simply a product of vectors whose elements are all equal to the variance added on the branches leading to the children. Hence the inverse of an individual blocks of **C**_*p*_ corresponding to child *c* is

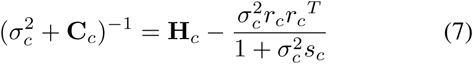

where **H**_*c*_ is the inverse of **C**_*c*_, 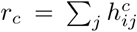 is the row or column sum of the symmetric matrix **H**_*c*_ and *s*_*c*_ is the sum of all elements of **H**_*c*_. This recursive update can be computed in 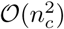 operations where *n*_*c*_ is number of leaves of the child *c*. In case the child node *c*_*i*_ is a terminal node, the respective block diagonal is the scalar 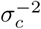. By traversing the tree from its leaves to root, each internal node can be dressed with a covariance matrix of its leaf nodes until we obtain the full covariance matrix between all leaves at the root. For a balanced tree, there are *𝒪*(log *n*) levels in the tree and the computational cost is dominated by the last operation at the root. Hence the overall computational complexity is *𝒪*(*n*^2^). For a maximally unbalanced tree, the *𝒪*(*n*^2^) operation has to be done *𝒪*(*n*) times such that the overall complexity in this worse case scenario is *𝒪*(*n*^3^).

## Speed and accuracy

To get a sense of the numerical accuracy and the empirical scaling of the time required for recursive matrix inversion, I generated trees and the associated covariance matrices from a Kingman coalescent process. I inverted these matrices using the matrix inversion routing in the `numpy.linalg` package and using the recursive algorithm outlined above. The recursive algorithm clearly scales more favorably with matrix size and is faster for matrices larger than 500 *×* 500. The simply python/numpy implementation is dominated by tree-traversal and memory allocation up to the matrix size of about 1000 *×* 1000 and scales as ∼ *n*^2^ afterwards. Optimized memory management would likely reduce runtime. The standard matrix inversion scales as ∼ *n*^3^ as expected.

The numerical accuracy of the recursive algorithm is lower than that of `numpy.linalg.inv.` Individual elements of **C** *·* **H** differ by about 10^*-*^13 from the identity matrix for matrices of rank *n* = 1000 while these residuals are about 5-fold lower for `numpy.linalg.inv.`

## Simultaneous calculation of H for every choice of root

Our discussion above singled out a specific node as the root of the tree. In general, however, the root is not known and to determine a plausible root *χ*^2^ is optimized with respect to the position of the root on the tree. Minimizing *χ*^2^ directly would require repeated calculation of **H** for various choices of the root. However, in analogy to message-passing approaches for inference on trees (Mézard and Montanari, 2009), the inverse covariance matrix can be calculated for every node of the tree in one post-order followed by one pre-order tree traversal.

Assume we have calculated the inverse ‘leaf-covariance matrices’ **H**_*p*_ for each internal node *p* in a post-order traversal as described above. We can now calculate the ‘outgroup-covariance matrices’, that is the covariance Ĉ of all outgroup leaves of node *p* relative to the base the branch *p* and its inverse Ĥ, see Fig. 2. This calculation is analogous to Eq. (7): Instead of the children of the node, the individual blocks are formed by the covariance matrices of the sister clades **C**_*s*_ and the outgroup-covariance matrix of the parent node **C**_*o*_, see Fig. 2.

With the two inverse covariance matrix **H**_*p*_ and Ĥ _*p*_ of tips on either end of branch *p*, we can now calculate the inverse covariance matrix for an arbitrary choice of root along the branch by again using Eq. (7) with the outgroup and in-group as children, see Fig. 2.

## Efficient linear regression with tree-like covariance matrices

If the only objective is to minimize *χ*^2^ with respect to model parameters, the matrix inverse is not necessary and the optimal model parameters can be determined in linear time as shown by (Ho *et al.*, 2014). Differentiating Eq. (4) with respect to *β* and *α,* one finds:

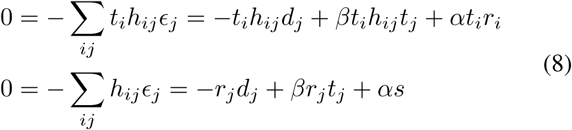

where repeated indices imply summation and *r*_*i*_ and *s* are the column sum and complete sum of *h*_*ij*_ as before. Straightforward algebra yields

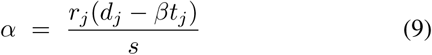

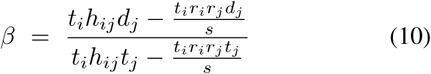

for the intercept and for the slope of the regression. The form of these expression is analogous to ordinary least squares regression 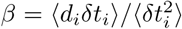 and *α* = 〈*d*_i_〉 – *β* 〈*t*_*i*_〉. The quantities ∑ _*i*_*t*_*i*_*r*_*i*_ and ∑ _*i*_*d*_*i*_*r*_*i*_ are essentially weighted averages of times and divergences with weights *r*_*i*_ (after dividing by *s* = ∑ _*i*_ *r*_*i*_). These weights *r*_*i*_ are lower if tip *i* is strongly correlated with several other tips *j*, see Fig. 1. The quantity *t*_*i*_*h*_*ij*_*t*_*j*_ and *t*_*i*_*h*_*ij*_*d*_*j*_, in turn, are analogs of second moments of time and distance. Since the matrix *h*_*ij*_ has block diagonal structure, these weighted sums can again be calculated recursively from the independent contributions of the children. The weighted sum at node *p* of sampling times obeys

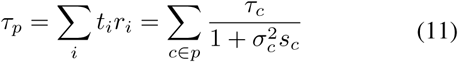

where *τ*_*c*_ is the equivalent sum of the child *c*. The only quantity that explicitly depends on the covariance matrix is the normalization *s*_*c*_ which obeys he recursive update 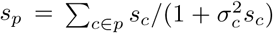.. These relations are analogous to those by Ho *et al.* (2014).

The divergence *d*_*pi*_ of tip *i*, one the other hand, depends on the reference node *p*. As the recursion moves up the tree towards the root, we can express the *d*_*pi*_ of tip *i* by the contribution of the branch *𝓁*_*c*_ leading to child node *c* and the *d*_*ci*_. The weighted sum therefore can be expressed as

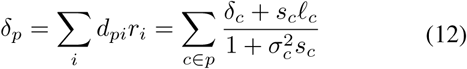

The second order quantities can be calculate analogously

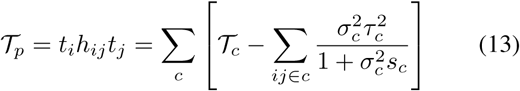

The situation is slightly more complicated for *t*_*i*_*h*_*ij*_*d*_*j*_ since as before the *d*_*j*_ change as we add another branch.

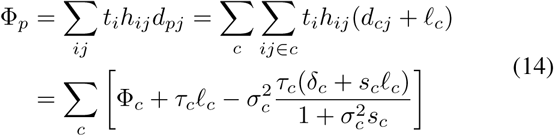

After evaluating these equations recursively in a post-order traversal, the quantities at the root node are the full weighted averages *τ* = ∑_*i*_ *r*_*i*_*t*_*i*_ etc. With these quantities in place, the optimal slope and intercept for a given choice of the root are given by

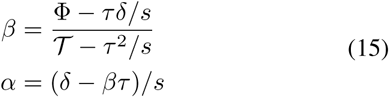

If the assumption that divergences are distributed as multivariate Gaussian is correct, the covariance matrix of the estimates is given by the inverse of the Hessian:

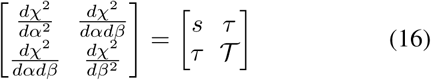

The covariance of the estimates can therefore be expressed as in terms of three scalar quantities that are already calculated.

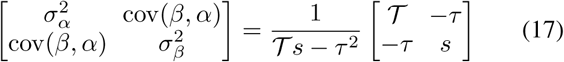

To obtain accurate estimates with tight confidence intervals, the weighted variance in tip dates *𝒯 /s - τ*^2^*/s*^2^ needs to be large, as one would intuitively expect.

### Optimizing the position of the root

The optimal root will rarely be an existing node, but will typically be an intermediate point on an existing branch at distance *𝓁x* and *𝓁*(1 *-x*) between two nodes, see Fig. 2.

In analogy to the calculations for the covariance matrix, we can calculate the quantities 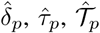, and 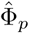 for all tips in the outgroup of branch *p*. If the variance 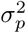 is linear in branch length, we can then calculate the weighted averages τ(*x*), *δ*(*x*), *𝒯*(*x*) and Φ(*x*) for any *x* by treating the outgroup and the in-group as two child nodes with branch length *x𝓁*_*p*_ and (1 *-x*)*𝓁*_*p*_. The resulting conditions are polynomial equations in *x* that don’t seem to have a convenient solution but are easily solved numerically.

### Practical issues

We have so far assumed that the variance contributions of the branches are known. However, these contributions are proportional to elapsed calendar time along the branch which is not known a priori. What we are given is branch length measured in the expected number of mutations which is proportional to time but fluctuates. Equating one with the other leads to systematic biases since less diverged parts of the tree will be associated with smaller variance such that they exert undue influence on the total estimate. This problem can be addressed with more complicated non-linear optimziation or by first estimating a time-scaled phylogeny using a rough rate estimate and then refining this estimate using branch length as measured in calendar time.

Fits of divergence time relationships with covariance aware methods can be sensitive to outliers that violate model assumptions. Since closely related tips are down-weighted due to their presumed covariation, outlier sequences have a larger weight in the covariance aware fit than in a simple least square fit. If the outliers are due to sequencing errors, culture adaptation, or similar artifacts, they should be removed.

Lastly, the accumulation of mutations is typically much more lumpy than predicted by a Poisson model. To improve stability of the rate estimation, it is advisable to include some extra variance in root-to-tip distance for each terminal branch in the model.

## Accounting for covariance reduces noise

To empirically investigate the improvement achieved when accounting for the covariance structure when estimating the rate of evolution and the time *T*_*M RCA*_ of the most recent common ancestor, I simulated evolution in population of size *N* = 100 under a neutral Wright-Fisher model using FFPopSim (Zanini and Neher, 2012). I sampled 200 individuals over a period of *T* = 2*N* generations, and recorded the true tree of the entire sample including the *T*_*M RCA*_. I reconstructed phylogenetic trees using IQ-TREE (Nguyen *et al.*, 2015) and estimated the substitution rate and *T*_*MRCA*_ using either root-to-tip vs time regression with naive least-squares or *χ*^2^ minimization. Accounting for the covariance structure dramatically reduced the noise in the estimates, see Fig. 4.

**FIG. 4.**
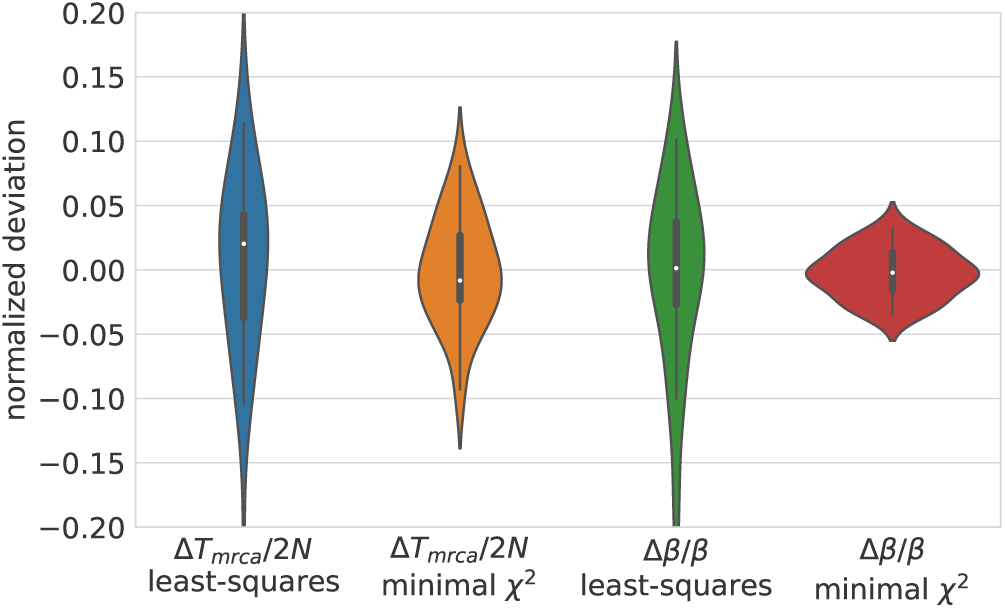
Accounting for the covariance structure of reduces variation in estimates of the *T*_*MCRA*_ and the evolutionary rate *β.* The figure compares results from simple least-squares regression with minimization of *χ*^2^ as defined in Eq. (4) on simulated data. Estimates that account for correlations in divergence among tips are more accurate.

## Implementation

The algorithm to invert covariance matrices and perform regression on trees is implemented as a class `TreeRegression` in TreeTime. Estimation of root-to-tip regression and clock rate estimation is exposed as a commandline tool `treetime clock`. The output of this command when applied to a set of influenza virus NA sequences is shown in Fig. 5. The corresponding data set is part of the collections of treetime examples and tutorials.

**FIG. 5.**
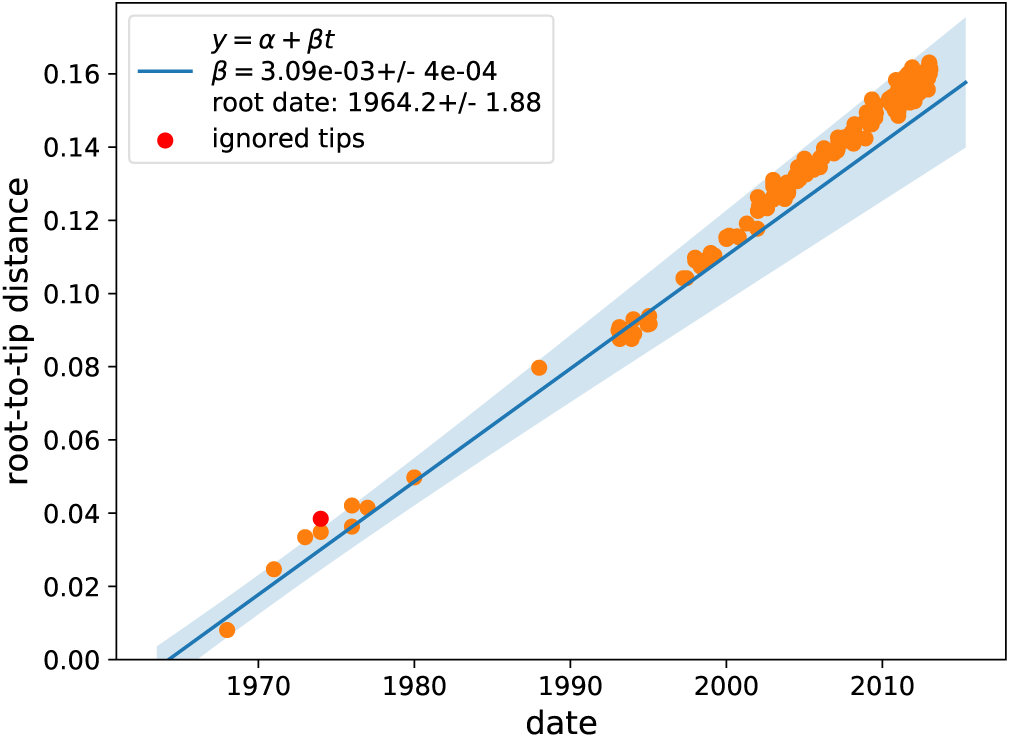
Optimized divergence vs time regression for 200 influenza A/H3N2 NA sequences as produced by treetime clock.

## Discussion

Phylogenetic correlations are a natural consequence of vertical descent and heritability and act as confounders in inference problems. But in the simple case of additive accumulation of variance along branches, these phylogenetic correlations can be readily taken into account. The recursive block like structure of the covariance matrix allows inversion using the Sherman-Morrison-Woodbury formula that has been used in a number of related problems (Hager, 1989). Ho *et al.* (2014) have shown that analogous algorithms hold for models in which variance does not accumulate linearly in time by saturates, e.g. in an Ornstein-Uhlenbeck process.

Here, I showed how the algorithm by Ho *et al.* (2014) can be used to simultaneously evaluate all possible choices for the root of the tree and estimate evolutionary rates. I included a slightly simplified and extended re-derivations of the recursive matrix inversion and linear model fit by Ho *et al.* (2014) here in the hope that they are helpful.

## Acknowledgements

I am very grateful to Vladimir Minin for pointing out the work by Ho and Ane.

## Appendix A: The value of the objective function

The objective function at optimal *α* and *β* is given by

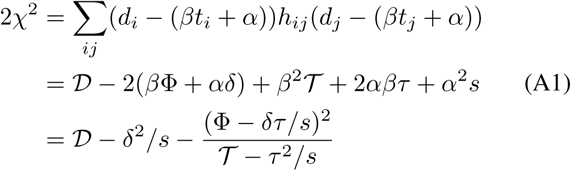

where *𝒟* is defined in analogy to *𝒯* and Φ

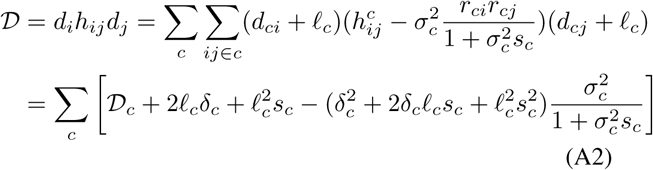

Here *𝒟*_*c*_ is the analogous quantity of the child and the full *𝒟* can be calculated recursively as before.

